# Need-selective gating of dopamine neuron cue responses by real and virtual hunger

**DOI:** 10.64898/2025.11.29.691326

**Authors:** Aphroditi A. Mamaligas, Joshua D. Berke, Anatol C. Kreitzer, Zachary A. Knight

## Abstract

The midbrain dopamine system is important for linking reward-predictive cues to learning and 20 motivation. Here we investigated how dopamine neuron responses to food and water cues are modulated by changes in internal state. We developed a flexible cued-approach task that allowed us to examine behavioral and neural responses to both food- and water-predictive cues within the same recording session. We found that overlapping subsets of dopamine neurons respond to food and water cues, but that the magnitude of these responses is gated in a need-25 specific way. Stimulation of hunger-promoting AgRP neurons amplified dopamine neuron responses to food cues, but not water cues, and the magnitude of these responses exceeded those observed in natural hunger. These findings indicate that changes in internal state modulate, in a need-appropriate way, the responses of a common set of dopamine neurons to environmental signals of food and water availability.

## INTRODUCTION

We find things that satisfy our internal needs rewarding. Thus, food tastes especially delicious when we are hungry. This connection between need and reward is essential to survival, both because it helps us learn about the value of different foods and fluids, and also because it helps 35 motivate our actions toward finding and consuming food and drink. Thus, it is important to establish the circuit mechanisms that connect need to reward in the brain.

The midbrain dopamine (DA) system is critical for linking rewards and reward-predictive cues to learning and motivation (Berke, 2018; Eshel et al., 2016; Salamone and Correa, 2012; Schultz 40 et al., 1997). DA neurons with cell bodies located in the ventral tegmental area (VTA) exhibit bursts of activity (∼20 Hz) in response to cues that predict the availability of food, water or other rewards (Eshel et al., 2016). This phasic activation – thought to function as a teaching signal that enables animals to learn associations between sensory stimuli and rewards in the environment (Schultz et al., 1997) – causes the release of DA in ventral striatum (also known as 45 nucleus accumbens, NAc (Day et al., 2007)). NAc DA release also plays a role in motivation,invigorating behavioral responses such as cued approach when the reward is expected (Blaiss and Janak, 2009; Hamid et al., 2016; Roitman et al., 2004; Wyvell and Berridge, 2000; Yun et al., 2004a, 2004b). Loss or blockade of DA reduces the willingness of animals to exert effort to obtain food or water, particularly when the amount of work required is high (Di Ciano et al.,50 2001; du Hoffmann and Nicola, 2014; Krause et al., 2010; Lex and Hauber, 2008; Stuber et al.,2008).

Given the importance of DA for motivation and learning, DA release should be modulated by internal state, so that an animal works harder to obtain goals that satisfy current needs.55 Consistent with this, it has been shown that DA release in response to food cues is greater in animals that are food deprived than those that are sated (Aitken et al., 2016; Cone et al., 2015; Konanur et al., 2020). Similarly, water deprivation amplifies the DA response to water cues (Fortin and Roitman, 2018; Hsu et al., 2020). This modulatory effect of internal state on the DA system has been proposed to be mediated by direct metabolic sensing by DA neurons (Abizaid60 et al., 2006; Cone et al., 2015) as well as by input from forebrain circuits that are specialized for monitoring energy and fluid balance. Particularly important nodes in this upstream circuitry include Agouti-related peptide (AgRP) expressing neurons in the arcuate nucleus of the hypothalamus (ARC), which are activated by food deprivation and promote hunger (Andermann and Lowell, 2017), and glutamatergic neurons in the subfornical organ (SFO), organum 65 vasculosum of the lamina terminalis (OVLT) and median preoptic nucleus (MnPO), which are activated by water deprivation and promote thirst (Grove and Knight, 2024).

However, it remains incompletely understood to what extent this state-dependent modulation is specific for cues that predict one particular reward over another, and, furthermore, how that specificity is encoded at the level of individual neurons. For this study, we developed a task in which we can monitor the behavioral and neural responses to different cues that predict either food or water availability, in the same animal and in the same session. We then used this task in conjunction with neural recordings and manipulations to address three questions: (1) Do food and water cues evoke responses in the same individual DA neurons, or in distinct DA subpopulations?, (2) How do changes in internal state modulate these response patterns? (3) What is the effect of AgRP neuron stimulation on these cue responses?

## RESULTS

### A task that measures responses to both food and water cues

Mice were placed in an operant chamber with two reward ports. On each trial, one of two tone cues (4.4 kHz or 12 kHz) was randomly presented; one tone predicted the availability of food, while the other predicted the availability of water. To obtain a reward, mice were required to nose-poke into the port that corresponded to the cue within 5 seconds of tone onset. Correct pokes (e.g. pokes into the food port following the food cue) immediately triggered delivery of that reward (either a 14 mg food pellet, or ∼5-10 µL of water; **Figure 1A**). Incorrect pokes resulted in a 5 second time out, signified by illumination of a white LED between the ports Mice (n=11) were subjected to mild restriction of both food and water and then trained in up to 8 training sessions (150 trials each). Mice were advanced from training to testing once they demonstrated mastery of the task, which we defined as less than 20% of trials incorrect (poke response in incorrect port during the 5 seconds of tone presentation) across a training session (49.6 ± 3.8% correctly completed trials on final training day; 6.0 ± 3.1% incorrectly completed trials on final training day). All animals trained were able to learn this task.

**FIgure 1.**
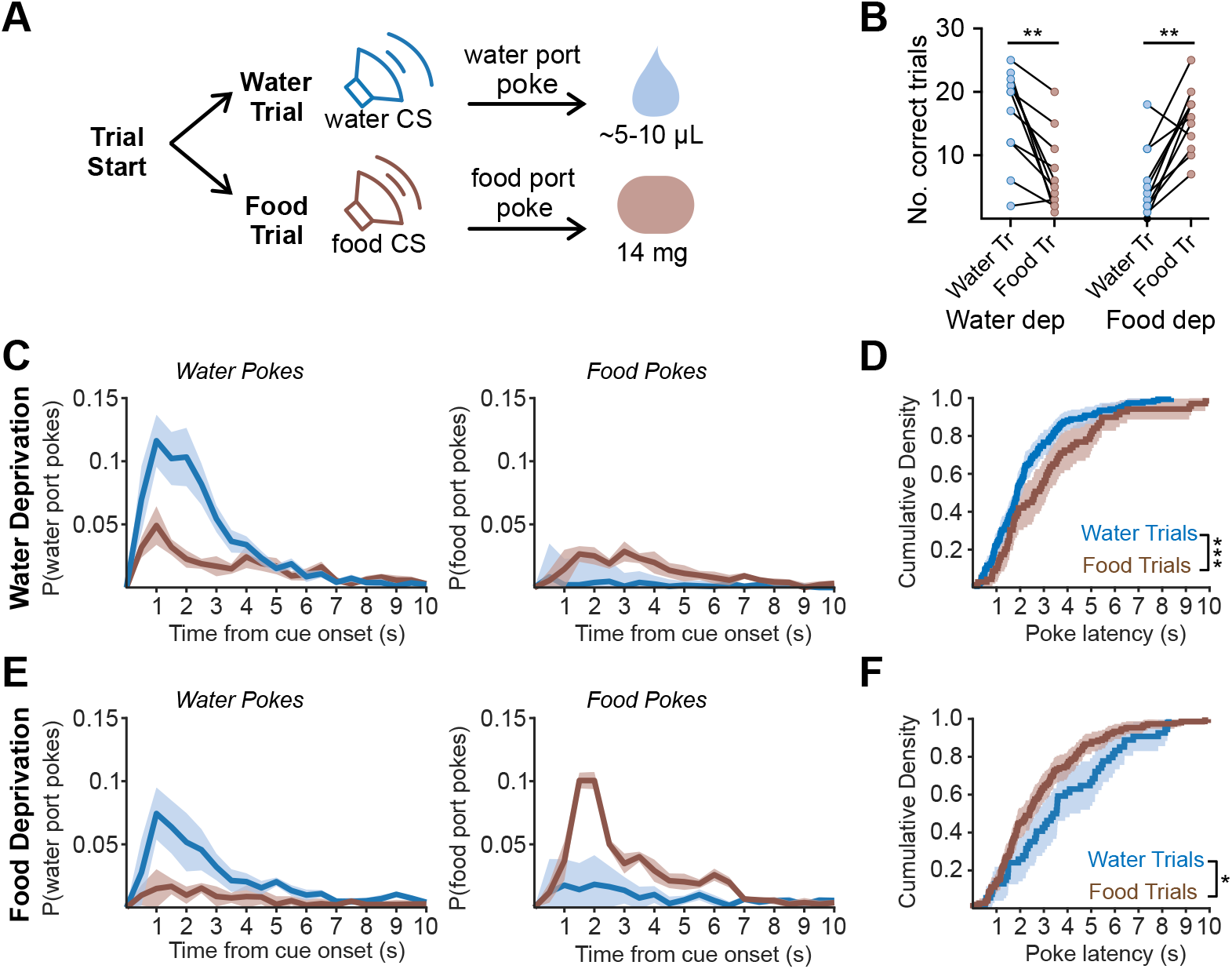
A task for measuring responses to food and water cues in the same session. A.) Schematic of task progression. During each trial, food or water-predictive tone will indicate ability for mice to poke for corresponding reward. B.) Mice correctly performed both food and water trials for a reward during food and water deprivation, with higher performance for the deprived reward. Wilcoxon matched pair signed rank. C.) Water deprived mice show a higher probability of entry into the water port during water trials (left graph) and the food port during food trials (right graph) following cue onset. D.) Distribution of poke latency for rewarded food and water trials in water deprived mice shows faster latency to enter the water port. Kolmogorov Smirnov test. E.) Food deprived mice show a higher probability of entry into the water port during water trials (left graph) and the food port during food trials (right graph) following cue onset. F.) Distribution of poke latency for rewarded food and water trials in food deprived mice shows faster latency to enter the food port. Kolmogorov Smirnov test. All summary values are displayed as mean ± SEM. * = p < 0.05, ** = p < 0.01, *** = p < 0.001.

We then measured performance in the task when mice had been overnight restricted for food or water, but not both, in order to determine how performance depends on need state. The test session consisted of 50 trials randomized between food and water trials. We found that, following food deprivation, mice were more likely to correctly perform food trials than water trials (68.4 ± 6% rewarded food trials vs 28.8 ± 6% rewarded water trials, n = 11, p = 0.001, Wilcoxon matched-pairs signed rank), whereas, following water deprivation, mice were more likely to perform more water trials than food trials (69.9 ± 6% rewarded water trials vs 28.3 ± 7% food trials, n = 11, p < 0.01, Wilcoxon matched-pairs signed rank) (**Figure 1B)**. Moreover, motivation (as assessed by latency to approach the food or water port) was correlated with the state of deprivation (**Figure 1D, F**). Under water deprivation, mice responded to the water cue by approaching the water port with shorter latency than the food port in response to the food cue (median water poke latency: 1.9 s, food poke latency: 2.9 s, p < 0.001, Kolmogorov Smirnov; **Figure 1D**). During food deprivation, the opposite was observed (water poke latency: 3.6 s, food poke latency: 2.4 s, p < 0.05, Kolmogorov Smirnov; **Figure 1F**). Thus, these data show that (1) mice can learn to differentiate between two auditory cues, one associated with food and one associated with water, and (2) their motivation to respond to these auditory cues is modulated by internal state.

### Bulk DA cue responses are selectively amplified by the corresponding need state

To investigate DA signals during this task, we targeted the calcium reporter GCaMP6f to DA neurons by a crossing a reporter line (Ai148) to mice expressing Cre from DA transporter locus (DAT^Cre^ mice). This resulted in GCaMP6f expression that overlapped with endogenous tyrosine hydroxylase (TH) in the midbrain (**Figure 2A**).

**Fig. 2.**
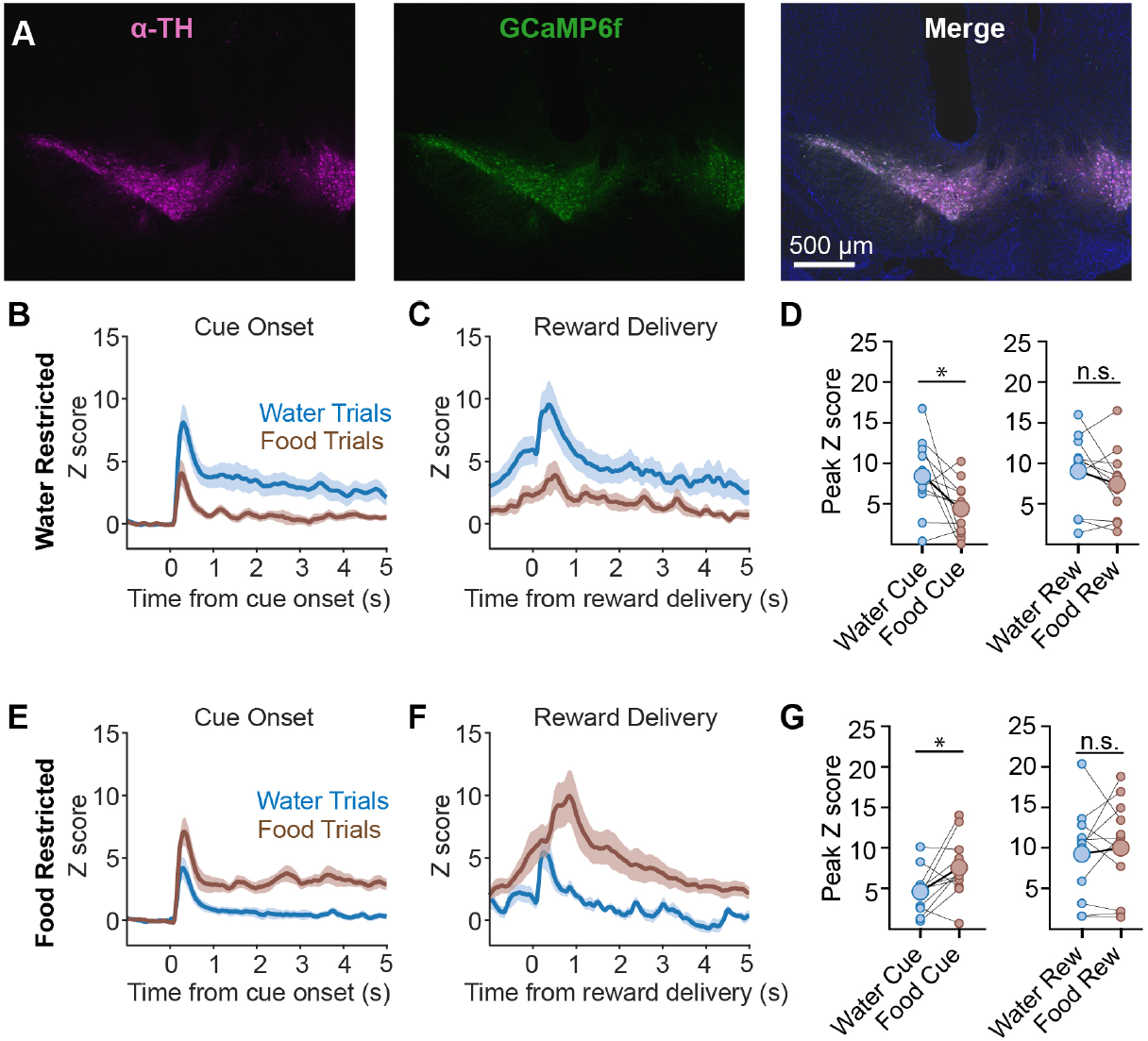
Dopamine neuron responses to food and water cues are modulated by need state. A.) Representative image of GCaMP6f expression in TH positive neurons in the VTA, as well as the fiber track of a photometry ferrule (merge; scale bar 500 µm). B.) Calcium signals aligned to cue onset in water deprived mice show a larger peak and extended activity in water trials than those in food trials C.) Calcium signals aligned to reward delivery in water deprived mice show a larger peak and overall activity in response to water trials. D.) Peak cue and reward-aligned GCaMP values in water deprived mice. Wilcoxon matched pair signed rank. E.) Calcium signals aligned to cue onset in food deprived mice show a larger peak and extended activity in food trials than those in water trials F.) Calcium signals aligned to reward delivery in food deprived mice show a larger peak and overall activity in response to food trials. G.) Peak cue and reward-aligned GCaMP values in food deprived mice. Wilcoxon matched pair signed rank.

We first implanted an optical fiber above the VTA for photometry measurements of dopamine neuron calcium activity. Mice were then trained in the task and tested on alternating days under either food or water restriction. As before, animals were subjected to 50 trials in which they were presented with either the food or water cue in a pseudorandom order. This revealed, first, that there were robust DA neuron responses to food- and water-predictive cues in both conditions (**Figure 2B-G**), indicating that cue responses are not strictly dependent on prior deprivation.However, the DA neuron responses to the deprived cue (i.e. the food cue in hungry mice and water cue in thirsty mice) were significantly larger than the responses to the non-deprived cue (water deprivation: water cue peak 8.40 ± 1.4 z, food cue peak 4.41 ± 1.0 z, n = 11, p < 0.05, Wilcoxon matched pair signed rank; food deprivation: water cue peak 4.62 ± 0.8 z, food cue peak 7.58 ± 1.1 z, n = 11, p < 0.05, Wilcoxon matched pair signed rank) (**Figure 2D,G**).Additionally, DA activity remained elevated for several seconds following presentation of the deprived cue, but not the non-deprived cue (mean signal between 1-3 s following cue presentation; water deprivation: water cue sustained signal 3.60 ± 0.9 z, food cue sustained signal 0.64 ± 0.3 z, n = 11, p < 0.05, Wilcoxon matched pair signed rank; food deprivation: water cue sustained signal: 0.60 ± 0.3 z, food cue sustained signal 3.02 ± 0.5 z, n = 11, p < 0.01,Wilcoxon matched pair signed rank) (**Figure 2B,E)**. This sustained response may be linked to consummatory behavior, because a peak in DA neuron responses was observed 0.5 to 1.0 sec after reward delivery (i.e. during consumption, **Figure 2C,F**). Taken together, these data indicate that (1) DA neurons respond to motivationally salient cues and (2) these responses are amplified for cues that predict satisfaction of a physiologic need.

### Food and water cues activate overlapping populations of DA neurons

Fiber photometry provides bulk measurements of neural activity but cannot reveal single-cell responses. While some studies have suggested that individual DA neurons are tuned to specific rewards or task variables (Engelhard et al., 2019; Grove et al., 2022), others have found broader responses (Lak et al., 2014; Millidge et al., 2024). It remains unclear to what degree there are specialized DA neurons that respond to food or water cues analogous to the specialized circuits in the forebrain that sense energy and fluid balance (Andermann and Lowell, 2017; Grove and Knight, 2024).

To investigate these possibilities, we targeted GCaMP6f to DA neurons and then implanted a gradient index (GRIN) lens over the VTA for single-cell calcium imaging of these cells (**Figure 3A**). Mice were trained in the task and then tested on alternating days in a state of either food or water deprivation while calcium dynamics were recorded by microendoscopy.

**Fig. 3.**
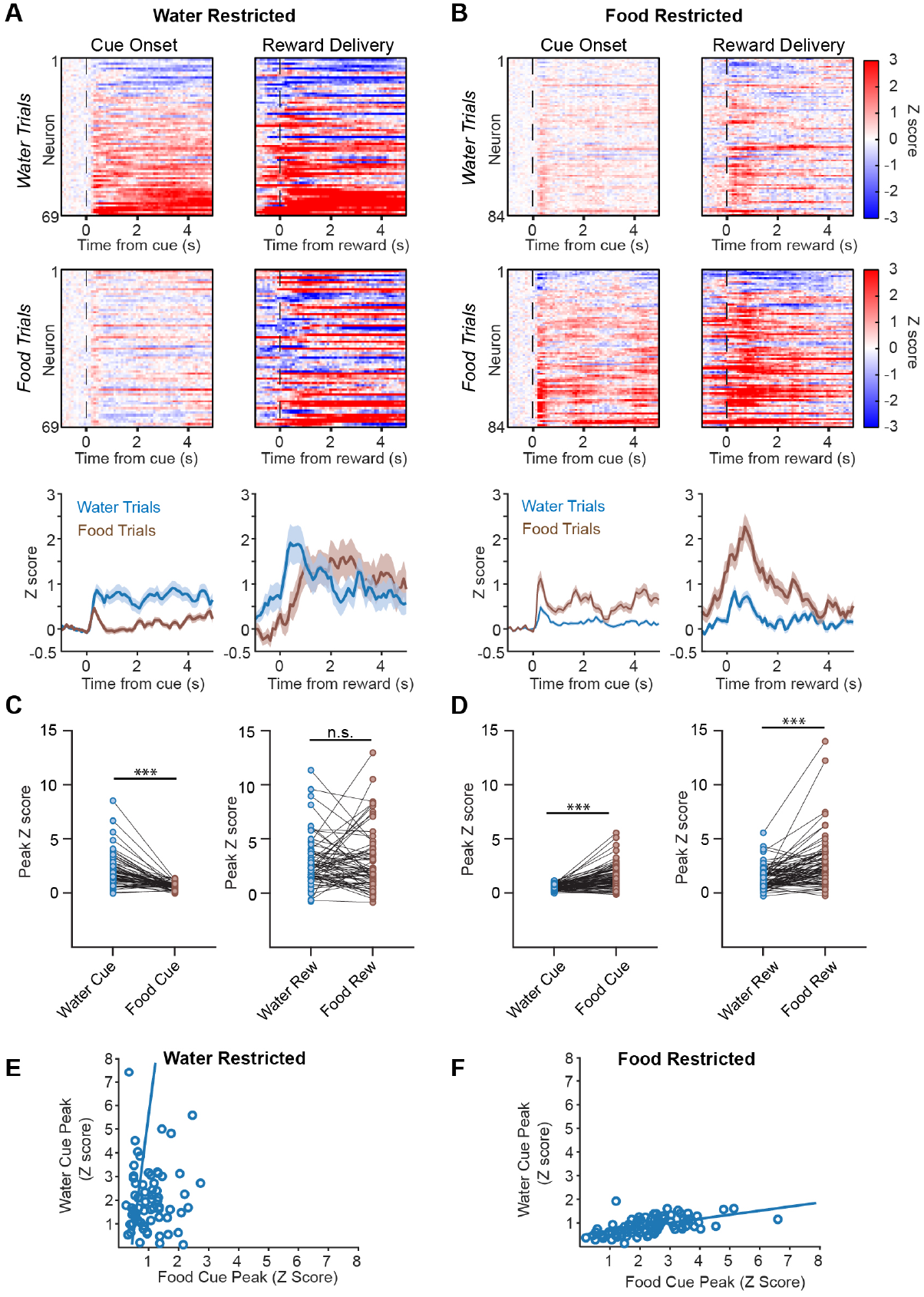
Responses of individual DA neurons to food and water cues. A.) Cue and reward-aligned single neuron calcium signals to food and water trials in water deprived mice show a stronger response to water trials (top: individual neuron heatmaps, bottom: summary data). B.) Cue and reward-aligned single neuron calcium signals to food and water trials in food deprived mice show a stronger response to food trials (top: individual neuron heatmaps, bottom: summary data). C.) Peak cue and reward-aligned z score calcium signals for each neuron in (A) for water deprived mice. Wilcoxon matched pair signed rank. D.) Peak cue and reward-aligned z score calcium signals for each neuron in (B) for food deprived mice. Wilcoxon matched pair signed rank.E.)Relationship between food and water cue amplitude in individual neurons for water deprived mice. F.)Relationship between food and water cue amplitude in individual neurons for food deprived mice. All summary values are displayed as mean± SEM. *** = p < 0.001.

Consistent with the results from fiber photometry, we found that a large proportion of individual DA neurons were activated by both food and water cues in each need state, but also that responses were greater and more sustained when the cue predicted satisfaction of a physiologic need (e.g. food cue in the food-deprived state; **Figure 3**). We found that for food deprived mice, peak activation for both food cue and reward aligned signals was greater than for water cue and reward signals (food cue aligned peak: 1.48 ± 0.12 z, water cue aligned peak: 0.51 ± 0.03 z, n = 84 neurons, p < 0.0001, Wilcoxon matched pair signed rank; food reward aligned peak: 2.63 ± 0.25 z, water reward aligned peak: 1.39 ± 0.11 z, n = 84 neurons, p < 0.0001, Wilcoxon matched pair signed rank) (**Figure 3B,D**). Meanwhile, in water deprived mice, while deprivation state strongly modulated cue aligned signals, this was not observed for reward aligned activity (water cue aligned peak: 2.02 ± 0.18 z, food cue aligned peak: 0.62 ± 0.04 z, n = 69 neurons, p < 0.0001, Wilcoxon matched pair signed rank; water reward aligned peak: 2.82 ± 0.27 z, food reward aligned peak: 2.85 ± 0.33 z, n = 69 neurons, p > 0.05, Wilcoxon matched pair signed rank) (**Figure 3A,C**). We also observed a second peak in DA neuron activity that occurred 0.5 to 1s after reward delivery and likely correlated with consummatory behavior. This peak was observed in food trials regardless of need state, whereas it was much smaller in water trials in the absence of dehydration. This may be due to differences in consummatory behavior during food and water deprivation.

We found no evidence to suggest that a high percentage of DA neurons are tuned to respond only to food or water cues. First, we found that most DA neurons responded to both food and water cues regardless of deprivation state, although the responses to the deprived cue were substantially larger. For example, in the water deprived state, 62% of neurons were activated by both food and water cues (out of a total of 69 neurons, p<0.05 by permutation test), whereas in the food deprived state, 72% of neurons were activated by both food and water cues (out of a total of 84 neurons, p<0.05 by permutation test). These percentages would presumably be higher if we were able to compare the cue responses of individual cells across different deprivation states, but we could not reliably align cells between days due to changes in the field of view.

Second, when we plotted the magnitude of the responses of individual DA neurons to food and water cues under a single deprivation state, we found that they were correlated (food deprivation: r^2^ = 0.30, p = 5.43e-8; water deprivation: r^2^ = 0.25, p = 5.66e-3). That is, neurons that responded strongly to food cues under food deprivation tended to also show stronger responses to water cues under food deprivation (**Figure 3F**) and vice versa for measurements under water deprivation **(Figure 3E**). If DA neurons were specialized for responding only to food and water cues, then we would expect these values to be uncorrelated. Together, these analyses indicate that most DA neurons in the VTA respond to both food and water cues.

### AgRP neuron stimulation selectively amplifies DA neuron responses to food cues

The enhanced DA neuron responses to food cues in food deprived animals could be mediated by direct metabolic sensing by VTA^DA^ neurons or the effect of upstream circuits. Given that AgRP neurons are important for many responses to food deprivation, we investigated the role of 200 these cells. We prepared mice for chemogenetic activation of AgRP neurons combined with microendoscope recordings of DA neuron activity. To do this we injected a Flp-dependent AAV expressing hM3Dq into the ARC, and a Cre-dependent AAV expressing GCaMP6f into the VTA of *Npy*^*Flp*^*;Dat*^*Cre*^ mice. In the same surgery, we implanted a GRIN lens above the VTA for calcium imaging. We validated the efficacy of hM3Dq expression in AgRP neurons of these animals by showing that IP injection of clozapine-N-oxide (CNO, 1 mg/kg) caused an increase in correct food trial performance in ad libitum fed mice (sated/saline: 21.9 ± 5 % food trials, sated/CNO: 64.8 ± 9 % food trials, n = 5 mice, p = 0.003, paired t-test).

We first tested mice that were non-deprived (ad libitum access to food and water) and were given a control injection of saline prior to testing. As expected, these animals exhibited only moderate DA neuron responses to food and water cues in our task (sated/saline; food cue aligned peak: 2.40 ± 0.49 z, water cue aligned peak: 1.93 ± 0.40 z, n = 33 neurons, p > 0.05, Wilcoxon matched pair signed rank), whereas responses to food and water delivery were somewhat larger (food reward aligned peak: 10.27 ± 2.6 z, water reward aligned peak: 3.60 ± 1.24 z, n = 19 neurons, p < 0.05, Wilcoxon matched pair signed rank) (**Figure 4A & 4D**).

**Figure 4.**
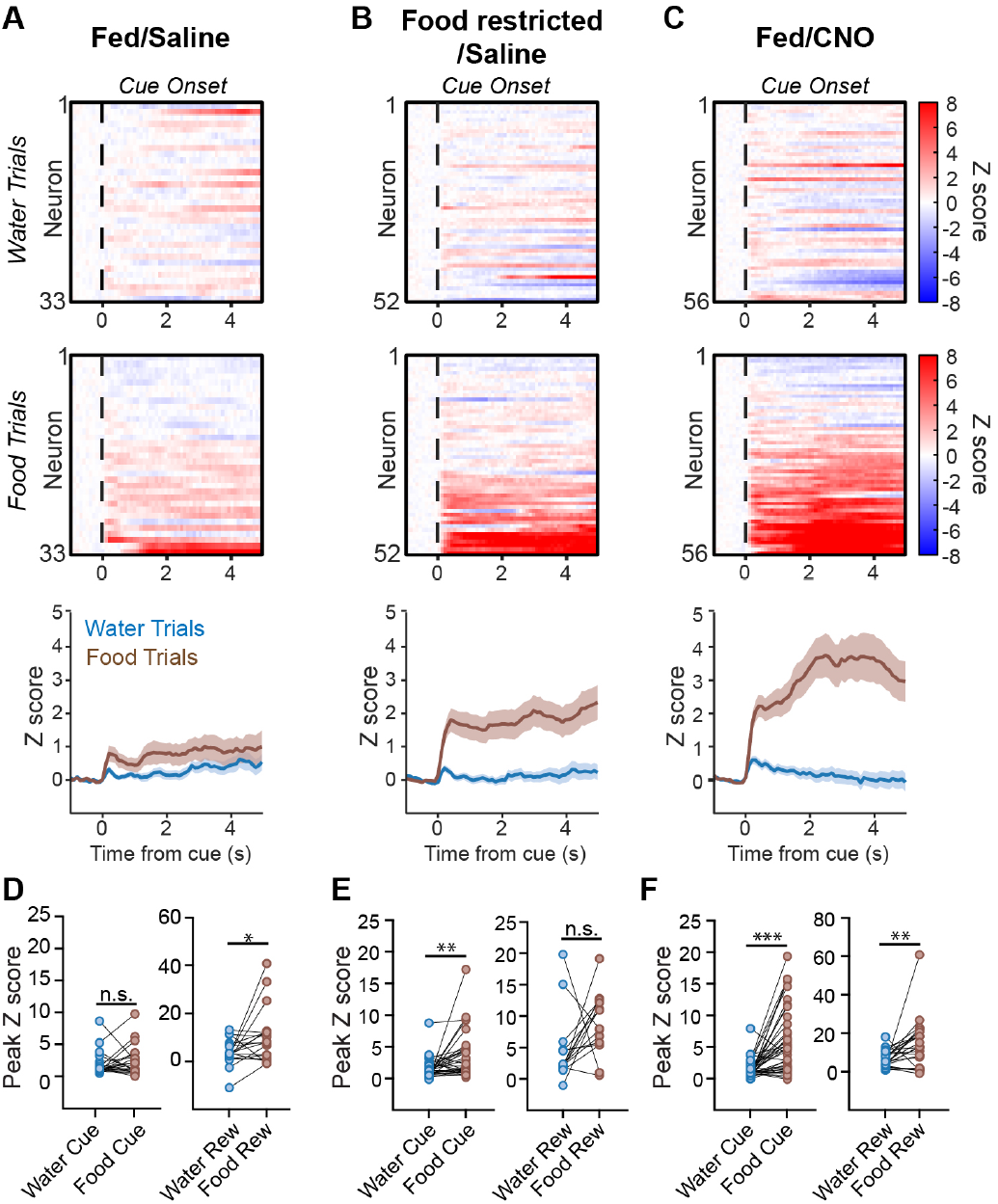
AgRP neuron activation selectively amplifies DA neuron responses to food cues. A.) Cue and reward-aligned single neuron calcium signals to food and water trials in sated mice injected with saline show weak activity of neurons overall (top: individual neuron heatmaps, bottom: summary data). B.) Cue and reward-aligned single neuron calcium signals to food and water trials in food deprived mice injected with saline show strong responses during food trials (top: individual neuron heatmaps, bottom: summary data). C.) Cue and reward-aligned single neuron calcium signals to food and water trials in sated mice injected with 1 mg/kg CNO show similarly strong responses during food trials that even overtook naturalistic hunger responses (top: individual neuron heatmaps, bottom: summary data). D.) Peak cue and reward-aligned z score calcium signals for each neuron in (A) for sated mice injected with saline. Wilcoxon matched pair signed rank. E.) Peak cue and reward-aligned z score calcium signals for each neuron in (B) for food deprived mice injected with saline. Wilcoxon matched pair signed rank. F.) Peak cue and reward-aligned z score calcium signals for each neuron in (C) for sated mice injected with 1 mg/kg CNO. Wilcoxon matched pair signed rank. All summary values are displayed as mean± SEM. * = p < 0.05, ** = p < 0.01, *** = p < 0.001.

Overnight food deprivation amplified the DA neuron responses to food cues but not water cues (food cue aligned peak: 3.62 ± 0.67 z, water cue aligned peak: 1.57 ± 0.30 z, n = 52 neurons, p < 0.01, Wilcoxon matched pair signed rank) **(Figure 4B**). This was due to an increase in the percentage of food cue responsive cells (sated/saline food responsive neurons: 45.4%, food deprived/saline food responsive neurons: 83.9%) whereas the mean strength of their activation did not significantly change (sated/saline food cue aligned peak: 2.40 ± 0.49 z, n = 33 neurons; food deprived/saline food cue aligned peak: 3.62 ± 0.67 z, n = 52 neurons; p = 0.11, Mann Whitney) (**Figure 4B, E**). Similar to earlier experiments (**Fig. 2 and 3**), we found that DA responses to food cues were sustained well beyond the cue itself, which likely reflects DA neuron responses to consummatory activity (sated/CNO sustained food cue signal: 3.94 ± 0.92 z, sated/CNO sustained water cue signal: 0.12 ± 0.24 z, p < 0.001, Wilcoxon matched pair signed rank) (**Figure 4B, E)**.

We next measured DA neuron responses in mice placed in a state of “virtual hunger” by AgRP neuron activation (CNO, 1 mg/kg, injected into sated mice 30 min before the start of the session). We found that AgRP neuron stimulation caused a striking increase in the DA neuron response to food cues, which even exceeded the response in food deprived animals (food deprived food cue aligned peak: 8.8 ± 1.27 z, n = 52; sated/CNO food cue aligned peak: 13.53 ±2.69 z, n = 56; p < 0.05, Mann Whitney) (**Figure 4C, E, F**). This was due to an increase in the percentage of responsive cells (sated/saline food cue responsive cells: 54.5%, n = 33 neurons; sated/CNO food cue responsive cells: 86.5%, n = 56 neurons) as well as the mean activation per cell (sated/saline food cue aligned peak: 2.40 ± 0.49 z, sated/CNO food cue aligned peak: 6.05 ± 0.86 z, p < 0.01, Mann Whitney). On the other hand, there was no effect of AgRP neuron activation on the response to water cues (sated/saline water cue aligned peak: 1.93 ± 0.40 z, sated/CNO water cue aligned peak: 1.45 ± 0.24 z, p > 0.05, Mann Whitney). This indicates that AgRP neuron activation is sufficient to recapitulate the effects of food deprivation on DA neuron activity, and that this modulation is highly specific for food versus water.

## DISCUSSION

Here we have investigated how DA neurons respond to food and water cues and how this is modulated by changes in internal state. We found that a broad population of DA neurons responds to auditory cues that predict the availability of either food or water, indicating that most DA cells are not tuned respond specifically to cues that satisfy one physiologic need over another. This is consistent with prior work showing that overlapping populations of DA neurons respond to exterosensory cues that predict different kinds of rewards (Lak et al., 2014; Millidge et al., 2024), although we are not aware of any studies that directly compare responses to food and water cues at the level of individual DA cells. Interestingly, DA neuron responses to post-ingestive food and water, which occur over longer timescales than responses to auditory cues, were shown to involve largely non-overlapping populations of DA neurons (Grove et al., 2022). This suggests that there may be different principles for how the DA system responds to exterosensory and interoceptive cues associated with ingestion.

Although food and water cues activate an overlapping population of DA neurons, we found that the strength of these cue responses is modulated by internal state in a need-specific way. Thus, food deprivation increases responses of individual DA neurons to food cues but not water cues (**Figure 3**). This is consistent with prior work showing, by fiber photometry, that DA neuron activity in response to a water-predictive auditory cue is potentiated by deprivation of water, but not by deprivation of food (Hsu et al., 2020).

AgRP neurons in the hypothalamic arcuate nucleus (ARC) are activated by food deprivation, and their artificial stimulation recapitulates the motivational and behavioral hallmarks of hunger (Andermann and Lowell, 2017). Therefore, AgRP neurons are a plausible candidate to link food deprivation to the modulation of DA neuron responses to food cues. Previous work has shown that DA release in NAc in response to food consumption is potentiated by stimulation of AgRP neurons (Alhadeff et al., 2019; Mazzone et al., 2020) and reduced by mutations that impair AgRP neuron function (Reichenbach et al., 2022). However, these studies did not clearly distinguish between DA neuron responses to food-predictive cues and DA neuron responses to reward delivery and consumption. They also did not measure how AgRP neuron stimulation modulates responses to cues that predict different kinds of rewards. We found that stimulation of AgRP neurons causes dramatic potentiation of DA neuron responses to an auditory food cue and, importantly, this potentiation is specific for cues that predict food versus water. This adds to other lines of evidence showing that the motivational effects of AgRP neuron stimulation are highly specific to food (Andermann and Lowell, 2017).

AgRP neurons track slow changes in energy balance through alterations in their tonic activity. Therefore, an important task is to explain how, and where in the brain, this tonic signal of energy balance is transformed into the modulation of phasic responses to food cues. The fact that most DA neurons do not seem to be specialized to respond only to food cues or only to water cues (**Figure 3**) implies that this modulation is unlikely to occur at the level of DA neuron cell bodies in the VTA, since any general change in excitability would presumably alter responses to both classes of cues. However, it is conceivable that axons innervating the VTA and relaying information about specific types of reward-predictive cues could be modulated in a need-specific way. AgRP neurons themselves do not project directly to the VTA, and the shortest path between the ARC and VTA likely involves an intermediate in the lateral hypothalamus (Fu et al., 2019). Alternatively, it is possible that modulation of cue responses by AgRP neurons occurs far upstream of the VTA at an early stage of sensory processing. For example, food deprivation has been shown to alter neural responses to food cues in postrhinal cortex, a visual association area, and this has been proposed to involve an AgRP→PVT→BLA→Ctx pathway (Burgess et al., 2018, 2016). An important task for the future will be to explain how the brain transforms these slow internal changes into the modulation of rapid cue responses, as this is central to understanding how need and reward are linked in the brain.

## Acknowledgements

This work was supported by National Institutes of Health grants R01-DK106399, R01-DK138127 and R01-DK145100 (to Z.A.K.), R01-NS1166226 (to Z.A.K., A.K., and J.B.), and F32-DA052283 (to A.M.). Z.A.K. is an Investigator of the Howard Hughes Medical Insitute.

## Declaration of Interests

The authors declare no competing interests.

## METHODS

### Animals

#### tereotactic surgery

All procedures were in accordance with protocols approved by the UCSF Institutional Animal Care and Use Committee. Mice were maintained on a 12/12 light/dark cycle and fed ad libitum. Experiments and surgeries were carried out during the light cycle. All surgeries were carried out in aseptic conditions while mice were anaesthetized with isoflurane (5% for induction, 0.5%– 1.5% afterward) in a manual stereotactic frame (Kopf). Buprenorphine HCl (0.1 mg kg-1, intraperitoneal injection) and Ketoprofen (5 mg kg-1, subcutaneous injection) were used for postoperative analgesia. Mice were allowed to recover for 14 - 42 days before experiments. For cell-type-specific expression of a calcium indicator in the VTA, we injected 500 nL of adeno-associated virus serotype 5 (AAV5) carrying GCaMP6f in a double-floxed inverted open reading frame under the control of the EF1a promoter (AAV5-EF1a-Flex-GCaMP6f; Addgene) into the VTA of DAT-Cre mice (Jackson Labs, strain #006660). This virus was injected unilaterally into the VTA of adult mice at coordinates AP -2.7, ML ±0.8 (counter-balanced to each side), and DV - 4.6, relative to bregma. Injections were performed using a pulled glass capillary with a Micro4 pump (WPI) at 100 nL/minute. In fiber photometry experiments, DAT-Cre mice were bred to the Ai148 GCaMP6f reporter line (Jackson Labs, strain #030328), resulting in expression of GCaMP6f in all DAT-expressing neurons throughout the brain.

For expression of the DREADD receptor hM3Dq in AgRP neurons of the arcuate nucleus, a Flp-dependent AAV construct expressing the hM3Dq receptor (AAV-DJ-fDIO-hM3Dq) was injected into the arcuate nucleus of DAT-Cre:NPY-Flp double mutant mice in addition to the GCaMP virus injected into the VTA as discussed above (NPY-Flp mice: Jackson Labs, strain# 030211). The hM3Dq virus was injected into the arcuate nucleus at coordinates AP -1.0, ML ±0.35, and DV -5.5.

To record calcium signals in vivo in freely-moving mice, animals were implanted with either a fiberoptic implant for fiber photometry or a gradient refractive index (GRIN) lens for single cell imaging into the VTA. Following the virus injections described above, the implants were inserted into the brain. For fiber photometry, a handmade 400 µm diameter, 5 mm fiber optic ferrule with a 0.38 numerical aperture was used. For single cell imaging, an integrated GRIN lens (Inscopix, Inc; 0.5 mm diameter, 6.1 mm length) was used. These implants were slowly lowered over 30 min to final coordinates AP -2.7, ML ±0.8, DV -4.5 using a motorized micromanipulator attached to the stereotaxic frame (Siskiyou). The implants were then secured in place with dental adhesive (C&B Metabond, Parkell), which was used to coat the surface of the skull, and a headcap was formed with dental acrylic (Ortho-Jet, Lang Dental).

#### Cued approach task

Mice were given food and water ad libitum before deprivation in preparation for training. Food or water restricted mice were placed in an operant box containing food and water ports. Two counterbalanced tones, differing only in frequency (4.4 kHz or 12 kHz), predicted either food or water availability in the respective port, and mice were trained to poke into the correct port within 5 seconds of tone onset to trigger reward delivery. Port entry was detected using custom-built ports with an infrared beam across the mouth of the port. The inter-trial interval between tone presentation was pseudo-random, following an exponential distribution between 5 and 30 seconds. Food pellets delivered were 14 mg standard chow pellets (Bio-Rad), and water reward was tap water (∼5-10 µL/reward). Water reward delivery was controlled by solenoids (NResearch) gating gravity-fed fluid delivery within the ports. Food delivery was controlled by a motorized food hopper (Med Associates). Incorrect pokes in which mice poked the opposing port relative to the tone were punished with a 5 second time-out, in which a white LED above the ports turned on and after which the inter-trial interval timer restarted. Similarly, this time-out was given following any inter-trial poke. During training, on this task mice were mildly restricted of both food and water such that they learned the task without bias to one reward over the other. Specifically, each mouse was given 90 minutes ad libitum access to water and 2g of food each day, at least one hour after training. Mice were trained until their error rate fell below a 20% criterion, but not for longer than 8 training days. Behavior experimental control, recording, and analysis were automated to remove possibility for experimenter bias. Age and sex of mice was balanced across cohorts, and littermate controls were used in each experiment where applicable. Behavior was controlled and recorded using MBED microcontrollers (Adafruit) interfaced to computers running StateScript behavior control software (SpikeGadgets). For a subset of experiments in animals expressing the DREADD construct hM3Dq, animals were injected with clozapine-N-oxide (Tocris, 1 mg/kg) or saline 30 minutes prior to beginning the task.

#### Histology

Animals were euthanized with a lethal dose of ketamine and xylazine (400 mg ketamine plus 20 mg xylazine per kilogram of body weight, i.p.) and transcardially perfused with PBS, followed by 4% paraformaldehyde (PFA). Following perfusion, brains were transferred into 4% PFA for 16 - 24 h and then moved to a 30% sucrose solution in PBS for 2 - 3 d (all at 4 C). Brains were then frozen and cut into 30 mm coronal sections with a sliding microtome (Leica Microsystems, model SM2000R) equipped with a freezing stage (Physitemp) and mounted on slides.

Slides were blocked for 1 hour in 10% Normal Donkey Serum (NDS) in 0.5% PBST then incubated overnight in primary antibody (1:500 rabbit anti-tyrosine hydroxylase, Pel-Freez P-40101), 3% NDS in 0.5% PBST. The following day, they were washed 3 times for 10 minutes each in 0.5% PBST and incubated for 2 hours in secondary antibody (1:750 donkey anti-rabbit 647, Invitrogen A-31573), 3% NDS in 0.5% PBST and 1:2000 DAPI. After this, slides were washed for 10 minutes in 0.5% PBST and 2 more 10 min periods with 1:1 PBS. Slides were then washed with 0.05% lithium carbonate and alcohol, rinsed with diH2O, and coverslipped with Cytoseal 60. Slides were scanned on a 6D high throughput microscope (Nikon, USA), globally gamma-adjusted to reduce background, and pseudocolored using freely available Fiji software.

#### In vivo imaging and analysis

Calcium imaging was performed using a head-mounted, microendoscope-based microscope to image through a chronically implanted GRIN lens placed above dorsal striatum (Inscopix, Inc). Blue light (435-460 nm) for excitation of the GCaMP6f calcium indicator was delivered at 0.5 mW/mm for continuous recording throughout the session. GCaMP6f emission signal (490-540 nm) was acquired continuously at 10 Hz and spatially down-sampled prior to motion correction. The motion-corrected video was then processed using Constrained Non-negative Matrix Factorization (CNMF) to isolate signals from individual neurons (Zhou et al., 2018). As was evident from visual inspection of the raw video, and similar to signals described in striatum (Barbera et al., 2016) and other brain regions (Jennings et al., 2015; Ziv et al., 2013), the individual neuron calcium signals resulting from this analysis were punctuated by brief, discrete, positive transients. These transients were detected post hoc using a threshold of 5 times the standard deviation of the full signal.

#### Fiber photometry

*In vivo* fiber photometry calcium data was acquired using a custom-built photometry system. An RZ5P fiber photometry processor (TDT) and Synapse software (TDT) were used to control LED output and acquire the photometry signal. Using this system, two LEDs were used to control GCaMP and isosbestic excitation (470 nm and 405 nm, respectively, Thorlabs). LEDs were sinusoidally modulated at 211 Hz (470 nm) and 531 Hz (405 nm) and entered a four-port fluorescence mini cube (Doric Lenses). The combined output was coupled to a fiber-optic patch cord (400 μm, 0.48 NA, Thorlabs), which then mated to the fiberoptic cannula in the mouse brain. The emitted light was collected onto a visible femtowatt photoreceiver module (AC low, Newport) and sampled at 60 Hz. Photometry data was then extracted via proprietary TDT software using MATLAB (MathWorks).

#### Statistics

Data are shown as mean ± SEM. Statistical significance was determined using Pearson correlation, Wilcoxon match-pairs signed rank test, Mann-Whitney test, Kolmogorov-Smirnov test, or Permutation test, where appropriate (MATLAB, MathWorks; Prism, GraphPad).

## Notes

### Competing Interest Statement

The authors have declared no competing interest.

